# SARS-CoV-2 infection, neuropathogenesis and transmission among deer mice: Implications for reverse zoonosis to New World rodents

**DOI:** 10.1101/2020.08.07.241810

**Authors:** Anna Fagre, Juliette Lewis, Miles Eckley, Shijun Zhan, Savannah M Rocha, Nicole R Sexton, Bradly Burke, Brian Geiss, Olve Peersen, Rebekah Kading, Joel Rovnak, Gregory D Ebel, Ronald B Tjalkens, Tawfik Aboellail, Tony Schountz

## Abstract

Coronavirus disease-19 (COVID-19) emerged in November, 2019 in China and rapidly became pandemic. As with other coronaviruses, a preponderance of evidence suggests the virus originated in horseshoe bats (*Rhinolophus* spp.) and likely underwent a recombination event in an intermediate host prior to entry into human populations. A significant concern is that SARS-CoV-2 could become established in secondary reservoir hosts outside of Asia. To assess this potential, we challenged deer mice (*Peromyscus maniculatus*) with SARS-CoV-2 and found robust virus replication in the upper respiratory tract, lungs and intestines, with detectable viral RNA for up to 21 days in oral swabs and 14 days in lungs. Virus entry into the brain also occurred, likely via gustatory-olfactory-trigeminal pathway with eventual compromise to the blood brain barrier. Despite this, no conspicuous signs of disease were observed and no deer mice succumbed to infection. Expression of several innate immune response genes were elevated in the lungs, notably IFNα, Cxcl10, Oas2, Tbk1 and Pycard. Elevated CD4 and CD8β expression in the lungs was concomitant with Tbx21, IFNγ and IL-21 expression, suggesting a type I inflammatory immune response. Contact transmission occurred from infected to naive deer mice through two passages, showing sustained natural transmission. In the second deer mouse passage, an insertion of 4 amino acids occurred to fixation in the N-terminal domain of the spike protein that is predicted to form a solvent-accessible loop. Subsequent examination of the source virus from BEI Resources indicated the mutation was present at very low levels, demonstrating potent purifying selection for the insert during in vivo passage. Collectively, this work has determined that deer mice are a suitable animal model for the study of SARS-CoV-2 pathogenesis, and that they have the potential to serve as secondary reservoir hosts that could lead to periodic outbreaks of COVID-19 in North America.

---

Coronavirus disease-19 (COVID-19) emerged in late 2019 in China and the etiologic agent was identified as a novel betacoronavirus, severe acute respiratory syndrome coronavirus-2 (SARS-CoV-2) (1-3). Both SARS-CoV and SARS-CoV-2 use angiotensin converting enzyme 2 (ACE2) as a cellular entry receptor in humans (2, 4). The virus likely originated from insectivorous horseshoe bats and may have undergone recombination in the receptor binding domain via an intermediate host prior to its spillover into humans (2, 5). To date, only a few mammalian species have been identified as susceptible, including Syrian hamsters, cynomolgus macaques, ferrets, felines, mink, Egyptian rousette bats and canines (6-12). Human ACE2-transgenic mice are also susceptible, unlike wildtype laboratory mice and rats (13).

A significant concern is the potential for spillback of SARS-CoV-2 into native wildlife species that could allow the virus to become endemic by the establishment of secondary reservoir hosts outside of Asia (14). Middle East respiratory syndrome coronavirus (MERS-CoV) likely transmitted from bats to dromedary camels in North Africa and established a secondary reservoir that accounts for most human outbreaks of MERS which repeatedly occur each year (15, 16). A recent report identified 20 important contact residues in human ACE2 for SARS-CoV-2 spike protein binding and suggested members of Bovidae and Cricetidae families may also be susceptible (17). Experimental challenge of Syrian hamsters, a cricetid rodent whose ACE2 shares 18 of these 20 critical residues, resulted in moderate, nonlethal pulmonary disease resembling human COVID-19 but without mortality (6, 7, 12).

Peromyscine rodents are also members of Cricetidae (subfamily Neotominae, genus *Peromyscus*) and are distributed throughout North America. There are 56 recognized species in the genus and some serve as reservoir hosts for diverse zoonotic agents, including hantaviruses, *Borrelia burgdorferi, Babesia* spp., and Powassan virus. Deer mice (*P. maniculatus*) are the most studied and abundant mammals in North America (18). The ACE2 receptor of deer mice shares 17 of the 20 critical residues for SARS-CoV-2 binding (Table S1), suggesting that deer mice may be susceptible to infection with SARS-CoV-2.

## Results

To test susceptibility, nine young adult deer mice 6 months of age of both sexes were intranasally challenged with 2×10^4^ TCID_50_ of SARS-CoV-2 and three were held as unchallenged controls. Three deer mice each were euthanized on days 3, 6 and 14 post inoculation (dpi) to assess infection, and one sham-inoculated deer mouse was euthanized on day 3 whereas the other two were euthanized on day 14. On days 3 and 6, gross substantial pulmonary consolidation and hemorrhage were observed in the cranial and middle portions of both lungs. Viral RNA (vRNA) was also detected in the lungs of these deer mice (Fig. 1A); however, by 14 dpi the lungs appeared healthy. Despite the resolution of the observed pathology during earlier time points, all three deer mice had low levels of viral RNA in their lungs at 14 dpi. Virus was also isolated from the lungs of each deer mouse euthanized on days 3 and 6, but not from deer mice euthanized on day 14 (data not shown). IgG to recombinant nucleoprotein was detected by ELISA only on day 14 (Fig. 1B; GMT=504), and low titer neutralizing antibody (GMT=25) was initially detected on day 6 that significantly increased on day 14 (GMT=160, Fig. 1C), suggesting neutralizing antibody on day 6 was likely IgM. Antibody to multiple viral antigens was detected in all 9 of the inoculated deer mice by western blot (Fig. S1), indicating an early antibody response.

**Figure 1.**
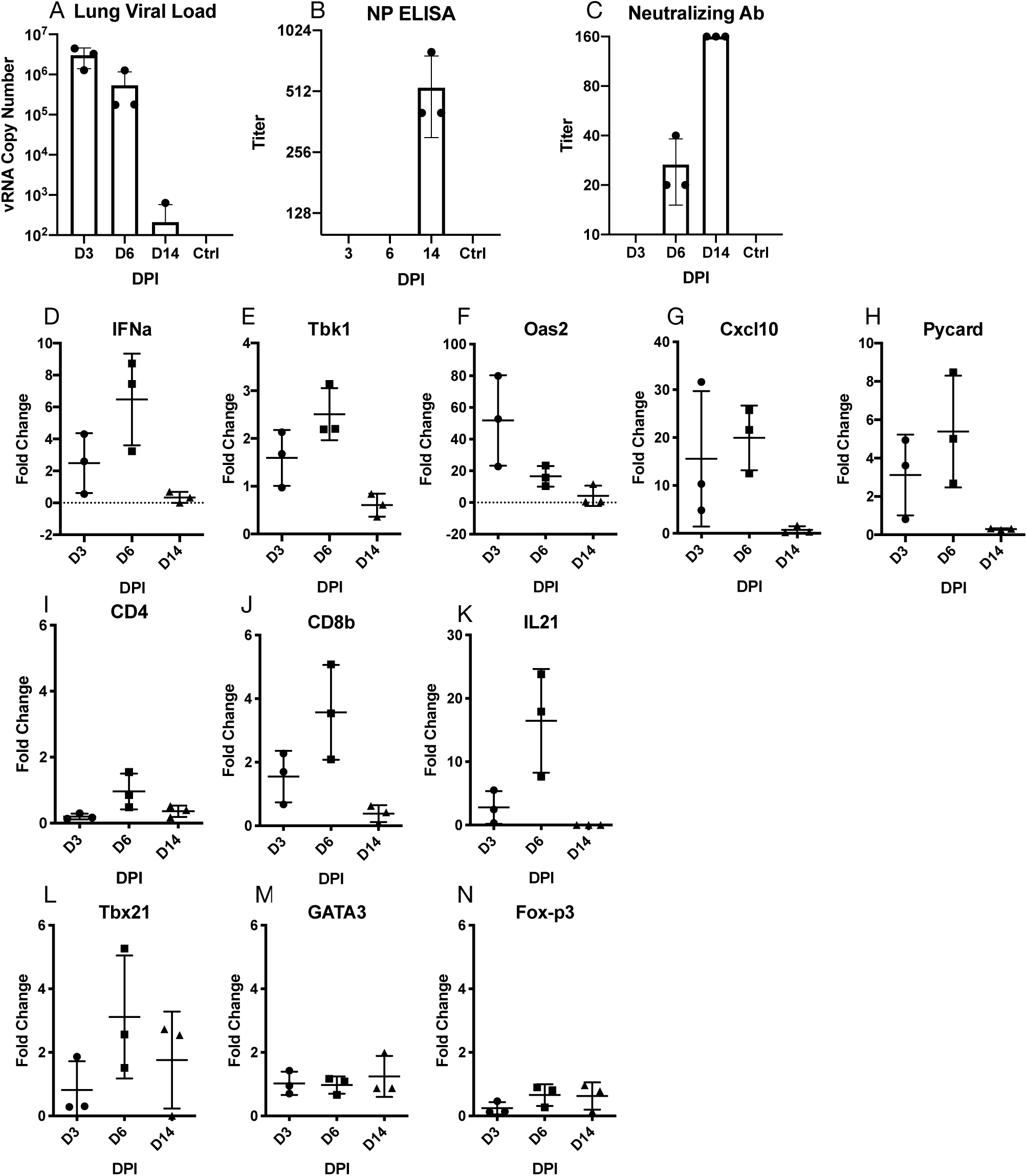
Infection and antibody response to SARS-CoV-2. (A) Viral RNA was detected in lungs of infected deer mice to day 14, (B) IgG antibodies were detected to nucleocapsid protein on day 14, (C) with neutralizing antibody detected on days 6 and 14. Each time point represents samples collected from 3 euthanized deer mice per group. Error bars represent the standard deviation of the mean (A) and geometric mean antibody titers (B, C). Virus was isolated from lung homogenates of day 3 and day 6 deer mice, but not day 14.

Examination of 41 immune response genes (19) (Table S2) identified several that were elevated in the lungs during infection. Five innate immune response genes (IFNα, Tbk1, Oas2, Cxcl10, Pycard) were substantially elevated during acute infection but then subsided by 14 dpi (Fig. 1D-H), indicating activation of antiviral defenses that declined as the virus was controlled. Expression of CD8β was substantially elevated on days 3 and 6, whereas CD4 expression was less elevated; both returned to nominal levels by day 14 (Fig. 1I-J). IFNγ expression was not detected in the lungs of the sham inoculated deer mice; however, it was detected at low levels in one day 3 deer mouse (Cq=35) and two day 6 deer mice (Cq=32, 35). IL-21 expression was greatest among the cytokines examined with up to 25-fold increase on day 6 (Fig. 1K). Considering the elevated expression of Tbx21 (T-bet), but not GATA3 or Foxp3 (Fig. 1L-N), the evidence suggests a pro-inflammatory type I immune response occurred in the lungs of deer mice.

Histologically, at 3 dpi, multifocal immunoreactivity was seen in occasional vibrissae with no associated inflammation in mystacial pad. Hemi skulls showed severe necrosuppurative to fibrinopurulent inflammation in nasal meatuses, maxillary sinuses and ethmoturbinates. Fibrinoid vascular necrosis obliterated the walls of medium-sized veins in ethmoturbinates. Severe ulceration and desquamation of the main olfactory epithelium (MOE) manifested in vomeronasal organ (VMO) and septum. Intense infiltrates of neutrophils surrounded and occasionally involved centrifugal afferent sensory branches of olfactory (CNI), ethmoidal, and maxillary nerves (Fig. 2A). Lingual fungiform and circumvallate papillae were surrounded by neutrophils, which clustered on necrotic gustatory buds. The tongue was mildly swollen due to dissecting edema and neutrophilic interstitial glossitis in all 3 deer mice at 3 dpi. Abundant viral antigen was present in the nasal passages spanning sustentacular, olfactory and basal cells of MOE in all 3 of the deer mice at 3 dpi (Fig. 2B) and less prominently in lingual mucosa, hard palate and oropharynx. Branches of chorda tympani, greater superficial petrosal and glossopharyngeal nerves (CNIX) were variably degenerate or minimally inflamed. Pterygopalatine, petrosal and lateral geniculate nuclei (in tympanic bulla) and trigeminal ganglion were multifocally degenerate and surrounded and/or infiltrated by neutrophils in 2 of the deer mice at 6 dpi. Virus antigen was present in the aforementioned ganglia at 3 and 6 days of infection (Fig. S2-S3). The glomerular layer of the main olfactory bulb (MOB) was spongiotic and immunoreactive at 3 dpi (Fig. 2C). Histioneutrophilic inflammation manifested in frontal lobe of the brain of one of the deer mice by 6 dpi (Fig. 2D-E). In inflamed areas of the MOB, viral antigen was also detected in cytoplasm of microglial and mitral cells (Fig. 2F-I). Less severe glial reactions and immunoreactivity were present multifocally within brain stem at the level of lateral sulcus nucleus (NTS), optic chiasm, hypothalamus, and thalamic parabrachial nucleus (PbN), ventral posteromedial nucleus (VPM) culminating in gustatory cortex. Retinal ganglionic and inner nuclear layers showed multifocal immunoreactivity (Fig. S4-S6). There were no significant histologic lesions in peripheral or central nervous systems at 14 dpi. Calvarial bone marrow showed multifocal cytoplasmic immunoreactivity in myeloid precursors, including megakaryocytes, at 6 dpi (Fig. S7). Inflammation and mucosal desquamation were mild to moderate in trachea and main stem bronchi showed mild to moderate immunoreactivity, respectively, at 3 dpi. Immunoreactivity was also detected in mononuclear and stellate, antigen presenting cells in reactive tracheobronchial and hilar lymph nodes at 6 dpi. Extensive histioneutrophilic and hemorrhagic bronchointerstitial pneumonia manifested at 3- and 6-days dpi along with leukocytoclastic vasculitis involving main and medium sized branches of the pulmonary artery along with marked peribronchiolar and perivascular lymphoid hyperplasia (Fig. S8). Perivascular mild neutrophilic infiltrates were present around main blood vessels at the base of the heart and around renal hilar vessels and nerves at 6 dpi (Fig. S8). Other visceral organs did not show significant pathologies. Occasional multinucleate cells of epithelial and histiocytic origin were scattered amongst inflammatory infiltrates. Inflammation significantly subsided at 14 dpi where lungs showed mild pulmonary fibrosis and residual pneumonia. Lamina proprial cellularity of the small intestine, particularly duodenum and ileum, moderately increased by 6 dpi and immunoreactivity was detected in crypt epithelium and mature enterocytes along with occasional submucosal macrophages (Fig. S9).

**Figure 2.**
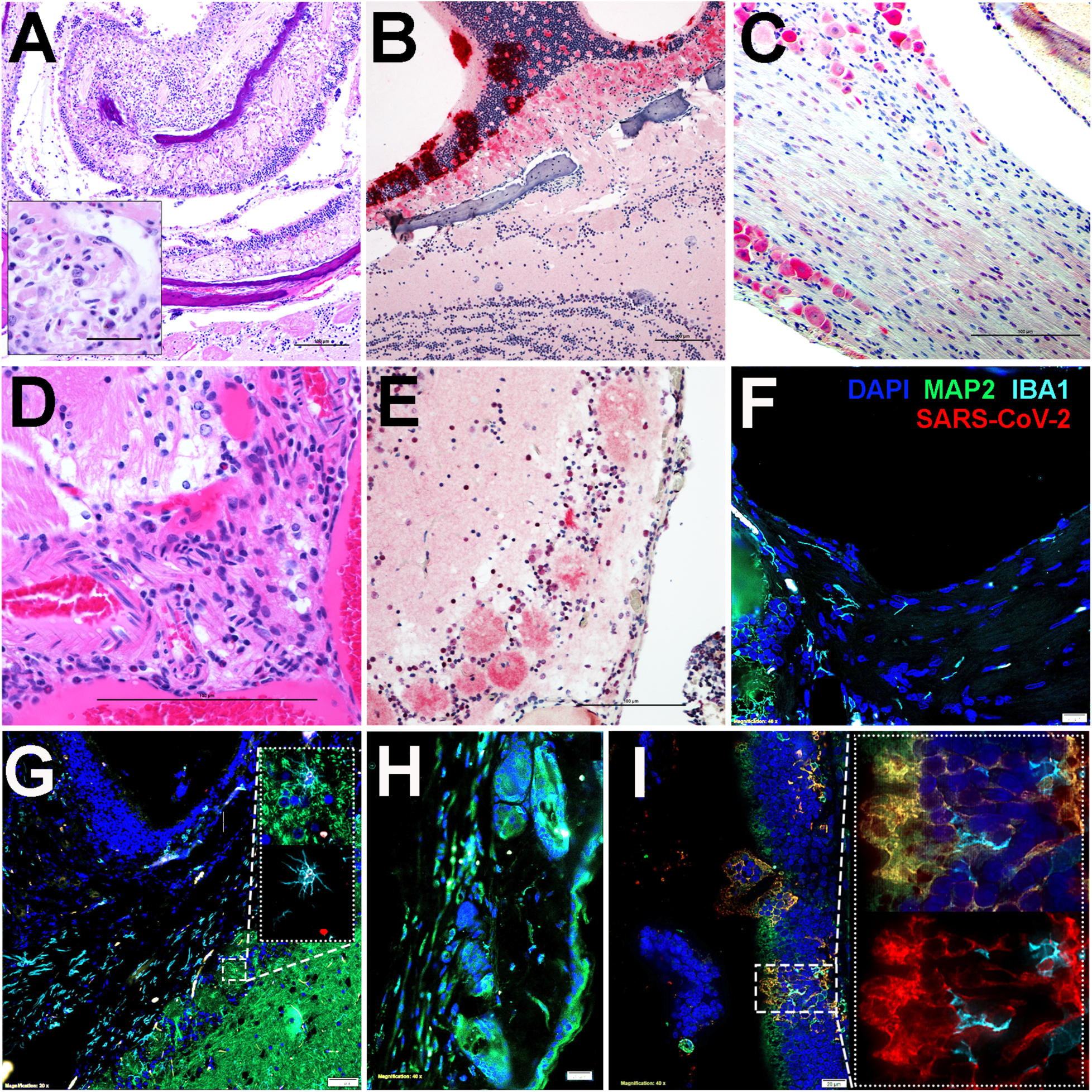
Histopathology and immunohistochemistry of SARS-CoV-2 in skull and brain of deer mice at 3- and 6-days post-infection. (A) Acute fibrinosuppurative and ulcerative sinusitis in ethmoturbinates with degeneration and inflammation of branches of olfactory, ethmoidal and maxillary sensory nerves (fibrinoid vascular necrosis – inset) 3 dpi. (B) Transmural SARS-CoV-2 immunoreactivity in MOE, 3dpi. (D) Prominent immunoreactivity in trigeminal ganglionic neurons with mild glial reaction at 6 dpi. Disruption of the BBB with associated histioneutrophilic meningoencephalitis of the MOB, 6 dpi. (E) Viral transmission to the glomerular layer of the MOB, 6 dpi. (F) Immunofluorescence imaging depicting entry of centrifugal afferents to the olfactory bulb at the cribriform plate in control mice 6 dpi after mock infection. Neurons were identified with antibodies against microtubule associated protein (MAP2, green), microglia with anti-ionized calcium-binding adaptor molecule 1 (IBA1, cyan), anti-SARS-CoV-2 (red) and nuclei were counterstained with DAPI (blue). (G) The presence of SARS-CoV-2 was detected by 6 dpi within neurons and microglia of the afferent nerves and in the glomerular layer of the olfactory bulb (100X high-magnification inset). Examination of the trigeminal nerve and ganglion in control (H) and infected (I) deer mice revealed neuroinvasion of SARS-CoV-2, with extensive co-localization of the virus within MAP2^+^ neurons proximal to activated microglia, 6 dpi (100X high-magnification inset). Bars equal 100 μm.

To assess transmission, 3 deer mice were intranasally inoculated with 2×10^4^ TCID_50_ of virus and the next day they were moved to a new cage containing 3 naive contact deer mice. Oral swabs, but not rectal swabs, of the three contact deer mice became vRNA^+^ on days 2, 5 and 5 post-contact (P1, Table 1). As they became vRNA^+^, the P1 deer mice were transferred to a third cage containing 2 additional naive contact deer mice (P2). The P2 deer mice had detectable vRNA from oral swabs 5 and 6 days after contact, demonstrating that sustained natural transmission can occur. Oral swabs of several deer mice that were vRNA^+^ became vRNA^-^ in subsequent days, only to become vRNA^+^ again (Table 1, Fig. 3). The inoculated and P1 deer mice were held until day 28 post contact, at which time all had antibodies detectable by western blot (Fig. S1). The P2 contact deer mice were euthanized on day 14 of the transmission study to provide a passage 2 viral stock. Each of the inoculated deer mice in the transmission study lost weight in the first 4-6 days before regaining weight, as did one passage 1 deer mouse (DM6) and both passage 2 deer mice (DM8, DM9) but to a lesser extent (Fig. 4A).

**Table 1.** Detection of vRNA in oral swabs.

| Study day | DM1 | DM2 | DM4 | DM5 | DM6 | DM7 | DM8 | DM9 |
| --- | --- | --- | --- | --- | --- | --- | --- | --- |
| **1** | + | + | + |  |  |  |  |  |
| **2** | + | + | + | Neg | Neg | Neg |  |  |
| **3** | + | + | + | + | Neg | Neg |  |  |
| **4** | + | + | + | + | Neg | Neg | Neg | Neg |
| **5** | + | + | + | + | Neg | Neg | Neg | Neg |
| **6** | + | + | + | + | + | + | Neg | Neg |
| **7** | + | + | + | + | + | + | Neg | Neg |
| **8** | + | + | + | + | + | + | + | Neg |
| **9** |  |  |  |  |  |  | Neg | + |
| **10** | + | + | Neg | + | + | Neg | + | + |
| **14** | + | + | + | Neg | Neg | + | Euth | Euth |
| **21** | Neg | + | + | Neg | Neg | + |  |  |
| **28** | Neg | Neg | Neg | Neg | Neg | Neg |  |  |
|  | Inoculated |  |  | Contact P1 |  |  | Contact P2 |  |

Gray boxes, no sample collected
+, vRNA detected by PCR
Neg, no vRNA detected by PCR
Euth, day of euthanasia of deer mice DM8 and DM9

**Figure 3.**
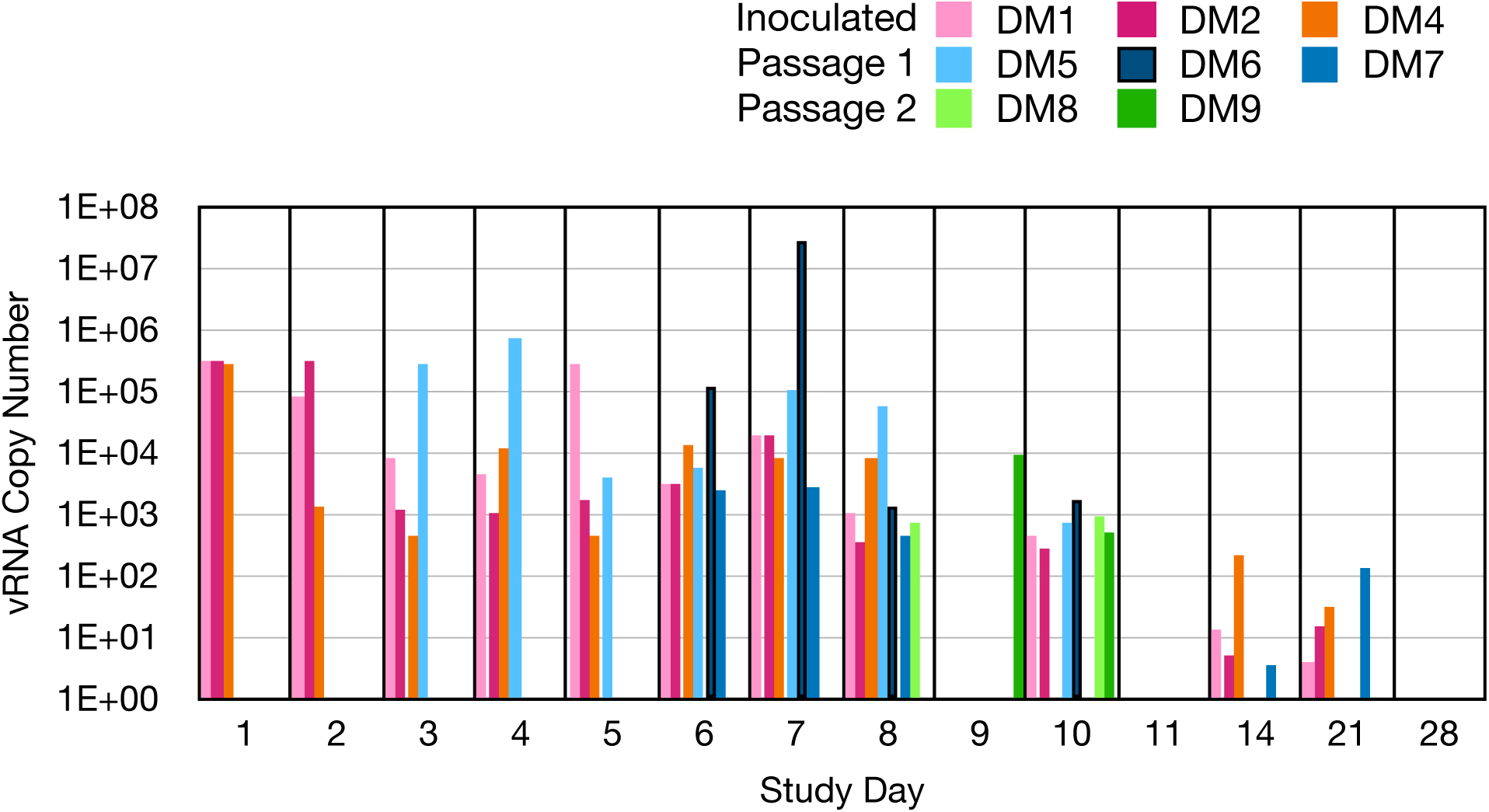
Detection of viral RNA in oral swabs of deer mice in the transmission study. Probe-based PCR was used to detect levels of E gene RNA and quantified against an E gene plasmid standard (Integrated DNA Technologies).

**Figure 4.**
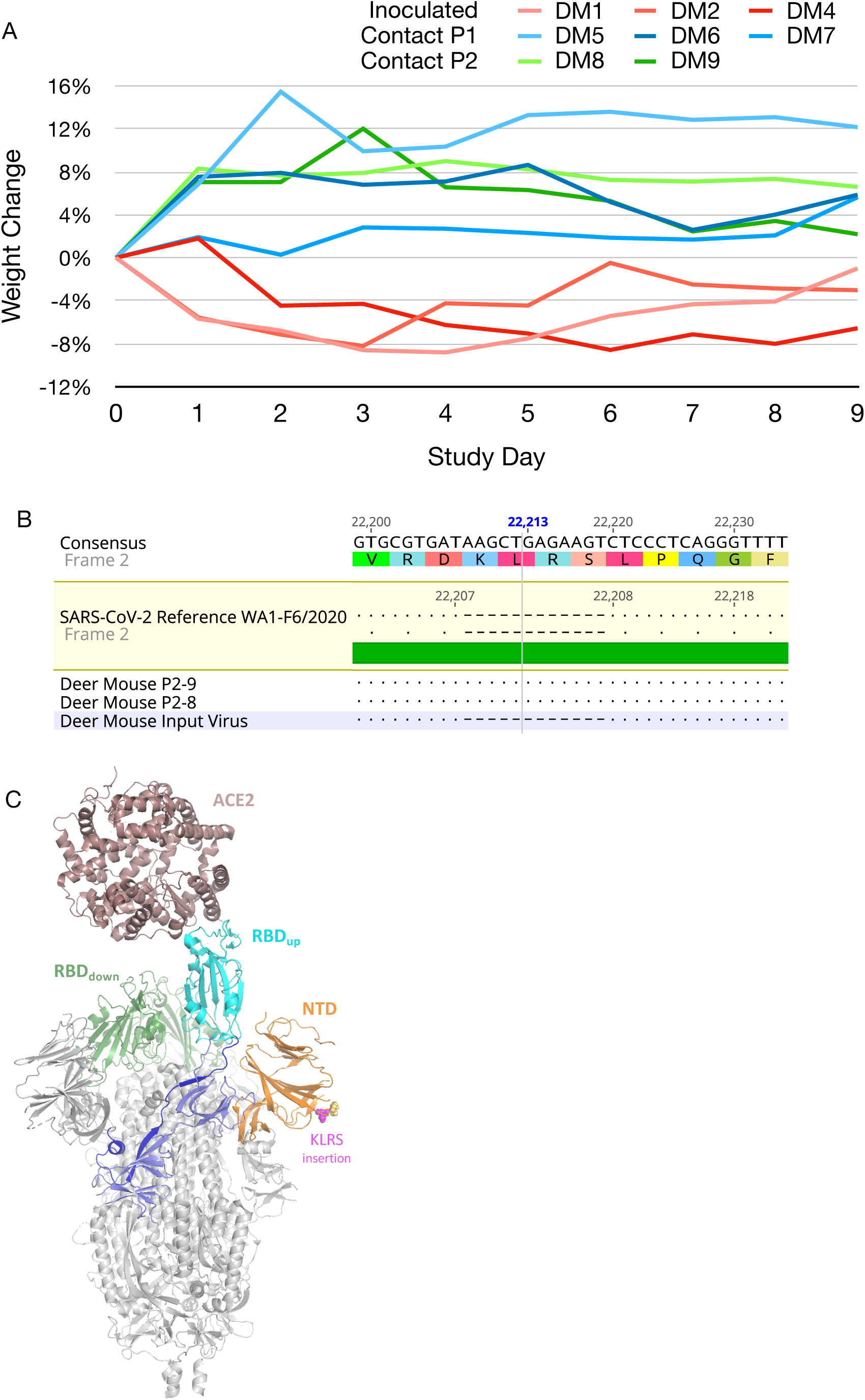
SARS-CoV-2 transmission in deer mice. (A) Transient weight loss occurred in some deer mice inoculated with SARS-CoV-2. Deer mice were weighed each day during the transmission study. All three inoculated deer mice (red lines) lost weight, one passage 1 deer mouse (DM6) and both passage 2 (DM8, DM9) deer mice. (B) Passage of SARS-CoV-2 led to fixation of a 4-residue insertion in the N-terminal domain. Oral swabs of both DM8 and DM9 passage 2 deer mice (P2-8, P2-9, respectively) were submitted to RNA-Seq and an insertion was detected in all reads spanning residues 216-219 (nt 22,208-22,209, hatches) of the spike protein, implying a strong purifying selection for the insert during passage. (C) Homology model of the spike protein-human ACE2 complex showing the location of the KLRS insertion observed in the P2 virus. Model was generated by threading the deer mouse P2 spike protein sequence into the EM structure of the open-state SARS-CoV-2 spike protein (PDB: 6VYB) (31) and colored as follows: Core (grey), N-terminal domain (NTD, orange) with KLRS insert as magenta spheres and native loop residues as yellow spheres, interdomain linker (blue), receptor binding domain (RBD, cyan for one up-conformation copy, green for two down-conformation copies), SD1/SD2 domains (light blue). The structure of the human ACE2 receptor (brown) was then superposed on the up-confirmation RBD using the crystal structure of the RBD-ACE2 complex (PDB: 6VW1) (32).

The viral RNA sequence was determined by NGS using oral swab samples from the two P2 deer mice and compared to the input sequence. A 4-residue insertion, KLRS, occurred in the N-terminal domain (NTD) of the spike protein, at residues 216-219 (nt 22,208-22,219) (Fig. 4B) and was observed in all reads (150 reads for DM8, 140 reads for DM9). Subsequent testing using a forward PCR primer that includes the 12 nt insertion showed the insertion occurred in all 17 infected deer mice in this study, implying a strong purifying selection. A structural model shows this insertion is located in a solvent-accessible loop that is distant from the RBD that interacts with ACE2 (Fig. 4C). The molecular origins of this selection are not yet know, but the surface location of the KLRS insert at a location away from the RBD may suggest interactions with an unknown co-receptor, and notably mouse hepatitis virus, also a coronavirus, uses the NTD as a receptor binding domain (20). It may be that the passage 2 deer mice were both infected by the passage 1 deer mouse DM5 (Table 1) that was introduced to the P2 cage when vRNA was initially detected in DM5 oral swabs 4 days prior to introduction of the other two passage 1 deer mice (DM6, DM7). However, the passage 2 deer mice did not become PCR^+^ until 2 days after DM6 and DM7 deer mice were added to the P2 cage, thus it cannot be conclusively determined which P1 deer mice transmitted the virus to the P2 deer mice.

Although the insertion was not detected in the input virus, which was sequenced at 8x coverage, conventional PCR using the forward primer containing the 12 nt insertion confirmed that the insert was present at low levels in the inoculum virus and at very low levels in the virus obtained from BEI Resources (Fig. S11). In addition, two synonymous mutations occurred in the Orf7 (codon 5, CUU > CUA) and nucleocapsid (codon 145, CAC > CAU) genes in the P2 virus. The weight loss of the three inoculated deer mice (Fig. 4A) could be reflective of a higher dose inoculum, or greater virulence of the wildtype virus that was abundant in the inoculum relative to the NTD insertion virus and will require further investigation. It remains unclear whether this insertion rose in frequency to near fixation due to bottlenecks during transmission and/or systemic spread of the virus, or to positive selection. Ongoing studies will address this issue directly.

## Discussion

The order Rodentia has more than 2,200 species and the divergence of Crictidae (hamsters, voles, lemmings, and New World mice and rats) and Muridae (Old World mice, rats and gerbils) families occurred about 18.5 mya (21). Cricetidae has more than 600 species, including at least 70 species in the North American subfamily Neotominae (21) that include deer mice and other peromyscines. Few cricetid ACE2 sequences are available but the 20 critical residues of ACE2 that interact with the RBD of SARS-CoV-2 (17) suggest other cricetid rodents are likely susceptible (Table S1). ACE2 residues are less human-like (15 of 20 residues) in the cricetid North American prairie vole (*Microtus ochrogaster*; subfamily Arvicolinae) and South American long-tailed pygmy rice rat (*Oligoryzomys longicaudatus*; subfamily Sigmodontinae), suggesting they are less likely to be susceptible. Neither the laboratory house mouse nor brown rat (family Muridae) are susceptible and each has 13 of the 20 critical ACE2 residues (17).

The respiratory pathology that occurs in deer mice demonstrates that they are a suitable small animal model for the study of SARS-CoV-2 disease. Elevated inflammatory immune signatures also suggested that a proinflammatory immune response may contribute to the pathogenesis of disease, which is also thought to be a significant contributor to COVID-19 morbidity and mortality (22). The elevated expression of CD8β, Tbx21 transcription factor and IL-21 suggest CTL infiltration and activation (23). We did not detect robust IFNγ or IL-6 expression, both of which are associated with fatal COVID-19 (24), and may account for why the deer mice did not have significant disease or death.

In addition, the presence of virus in the brain suggests a wide spectrum of neuropathologies could precipitate parasympathetic and sympathetic clinical symptoms such as xerostomia, epiphora, trigeminal neuralgia, confusion, and more consistently hyposmia/anosmia and hypogeusia/ageusia (25, 26). Bidirectional trans-neuronal dissemination in the medulla can result in rapid death if the more vital neighboring respiratory center becomes infected. SARS-CoV-2/deer mouse infection studies may clarify the complex relationship between sensory losses, chronic pain and compromise to the BBB and immune system of affected COVID-19 patients. If cervical ganglia are infected, autonomic dysfunction, and potential myogenic effect in addition to brain stem injury could lead to global brain ischemia due to electrographic myocardial injury and constriction of cervical and skull arterial blood supply (27). Collective pathology and immunohistochemistry results point to two different mechanisms that SARS-CoV-2 may utilize to enter the central nervous system. Similar to herpesviruses, early in the course of the disease at 3 dpi, the virus appeared to invade the brain in a retrograde axonal transmission along gustatory, olfactory and trigeminal pathways, bypassing the blood-brain barrier (BBB) (28).

Deer mice are among the most widely studied rodents in North America and are frequently collected by mammalogists conducting field work. The susceptibility of deer mice heightens concerns for reverse zoonosis whereby spillback of SARS-CoV-2 might occur that could lead to establishment of deer mice or other neotomid rodents as secondary reservoirs of SARS-CoV-2 in North America. Dromedary camels, which are secondary reservoirs of MERS-CoV-2, had detectable vRNA in oral swab samples for up to 35 days in an experimental infection study (29). All contact deer mice in this study became infected with detection of vRNA in oral swabs within 2 days of contact, and inoculated deer mice had detectable vRNA to 21 days, suggesting that sustained transmission among wild deer mice is possible. The susceptibility of two cricetid rodents, deer mice and Syrian hamsters, also raises the possibility that cricetid rodents could play roles in recombination events between coronaviruses. As a precedent, alphacoronaviruses have been detected in cricetid rodents in Europe, including bank voles (*Myodes glareolus*) and field voles (*Microtus agrestis*), and in this study each of the alphacoronaviruses possessed spike genes derived from betacoronaviruses (30). Considering that cricetid rodents are also found in Asia, it must be considered that they could have served as intermediate hosts for the recombination event that is thought to have occurred in SARS-CoV-2 prior to spillover into humans (5).

Deer mice and other peromyscine rodents have been used in biomedical research for many decades, including for their roles as reservoir hosts of zoonotic agents, and aging and diabetes studies (31), both of which are comorbidities associated with higher mortality rates in humans (32). They can live eight years in captivity, about four times longer than laboratory mice and Syrian hamsters, making them particularly suitable as a model organism to examine the effects of age and SARS-CoV-2 infection, and the durability of immunity to infection and vaccination. Moreover, as an outbred model, deer mice are more likely to reflect the diverse outcomes of infection observed in humans (e.g., neuropathology), in contrast to those that occur in highly inbred laboratory mice and Syrian hamsters (33).

## Methods

### Virus and Cells

SARS-CoV-2 (isolate 2019-nCoV/USA-WA1, NR52281) was provided by BEI Resources. Virus was passaged twice on Vero E6 cells (ATCC CRL-1586) in 2% FBS-DMEM containing 10,000 IU/ml penicillin and streptomycin at 37°C under 5% CO_2_ to generate stock virus used in these experiments. Virus was stored at −80°C. All work with infectious virus was performed at BSL-3 with approval from the Colorado State University Institutional Biosafety Committee.

### Animal Procedures

Deer mice (*Peromyscus maniculatus nebrascensis*) of both sexes and of 6 months of age were kindly provided by Dr. Ann Hawkinson at the University of Northern Colorado. This colony was established with deer mice captured near Whitewater, CO in 2000 (34). All animal work was approved by the Colorado State University Institutional Animal Care and Use Committee (protocol #993).

Deer mice were intranasally inoculated under inhalation isoflurane anesthesia with 2×10^4^ TCID_50_ SARS-CoV-2. On days 3, 6 and 14, three inoculated deer mice were euthanized, and one sham-inoculated deer mouse was euthanized on day 3 and the other two on day 14. Necropsies were performed for tissue RNA, virus isolation, and histopathology and immunohistochemistry.

To assess contact transmission, three deer mice were inoculated as described above. The next day, the deer mice were added to a cage with three naive contact deer mice (passage 1). Oral swabs were collected daily to determine infection status via qPCR. As the contact deer mice became PCR^+^ they were moved into another cage containing two additional naive contact deer mice (passage 2).

### Virus Detection

Swabs in virus transport medium were vortexed thoroughly and centrifuged to pellet cellular debris. RNA was extracted from swab supernatant using the QiaAmp Viral RNA kit (Qiagen) according to the manufacturer’s instructions. Tissues were homogenized and RNA extracted using RNeasy Mini kit (Qiagen) following manufacturer instructions. For detection of viral RNA, previously published assay (“one-step real-time RT-PCR E assay”) using the One-Step RT-qPCR kit (Roche Diagnostics). For each run, 3 µL dilutions of synthetic RNA standards were run in duplicate to quantify copy number in each sample. The Berlin E gene primer/probe/ plasmid standard (IDT Technologies) kit was used to quantify viral copy numbers (35).

### Serological Analysis

Truncated SARS-CoV-2 nucleocapsid (N) protein (AA133-416) was produced to reduce non-specific binding during antibody production (36, 37). A bacteria-codon optimized gBlock for the truncated protein was produced by IDT DNA Technologies and cloned into a pET28a bacterial expression with a C-terminal 6xHis tag using the NEBuilder Assembly Kit (New England Biolabs). Recombinant protein was expressed in BL21(DE3) pLysS *E. coli* and purified by nickel affinity and size exclusion chromatography essentially as previously described (38), with the exception of the use of 50 mM HEPES buffer (pH 7.4) and 500mM NaCl throughout purification to reduce aggregation. Purity and identity of purified nucleocapsid protein was determined by SDS-PAGE and mass spectrometry at the CSU Proteomics and Metabolomics Core, respectively.

ELISA was performed by coating plates with 200 ng/100 µl recombinant N protein diluted in PBS (pH 7.2) overnight then washed 3x with PBS. Plates were blocked with SuperBlock T20 (Thermo Scientific) for 30 minutes and washed. Heat inactivated serum samples (60°C 60 min) were diluted 1:100 in PBS, then serially diluted 2-fold and incubated with antigen for 1 hr at room temperature then washed 3x PBS-0.25% TWEEN20/3x PBS. Goat anti-*Peromyscus leucopus* IgG(H&L)-HRP conjugate (SeraCare) at 1:1,000 was incubated for 1 hour followed by PBS-TWEEN20/PBS washing. ABTS substrate (Thermo Fisher) was added and after 15 min absorbance (405 nm) recorded. The titers we determined by the reciprocal of the greatest dilution that was 0.200 OD above the mean of the negative control serum samples.

For western blot detection, infected Vero cell supernatants were subject to centrifugal filter concentration with molecular weight cutoff of 300 kDa (Pall Microsep 300K Omega). Enriched virus was inactivated with 2% sodium dodecyl sulfate (SDS) and protein concentration determined with a Pierce BCA protein assay kit (Thermo Scientific) according to manufacturer’s instructions. Eight micrograms of protein per lane were separated by 12% SDS polyacrylamide gel electrophoresis and blotted onto Immobilon-P nylon membranes (Millipore). After transfer, the blots were sectioned by lane, blocked, and individual lanes incubated with 1:100 dilutions of the indicated deer mouse sera or with mouse anti-nucleocapsid monoclonal antibody overnight. Primary antibodies from deer mice were detected with goat anti-*P. leucopus* IgG-HRP and developed with the TMB membrane peroxidase substrate system (3, 3 ‘, 5, 5’ - Tetramethylbenzidine, KPL). Images were scanned with a Visioneer One Touch 9420 scanner at a gamma value of 1.0, and all contrast adjustments were uniformly applied using Adobe Photoshop.

Serum neutralization was performed starting at 1:10 dilution with 2-fold dilution series. An equal volume of SARS-CoV (100 TCID_50_) was added (final dilution of 1:20) and incubated for 1 hr at 37°C. The mixture was plated on Vero E6 cells and scored for CPE after 6 days. The titer was reciprocal of the greatest dilution that conferred 100% protection.

### Immune Gene Expression Profiling

Deer moue immune gene expression profiling has been previously described (19). Briefly, primers (Table S2) for various immune genes were used to amplify cDNA collected from lungs using QuantiNova reverse transcription and SYBR Green I PCR kits (Qiagen). The ΔΔCq method was used with normalization within sample on GAPDH (ΔCq) and fold-change calculated for each gene against the means of the 3 sham inoculated control deer mice (ΔΔCq).

### Pathology

Three deer mice were humanely euthanized at 3-, 6- and 14 dpi and whole deer mice were freshly fixed in 10% neutral buffered formalin (10% NBF) after opening abdominal and thoracic cavities to allow for gross inspection and formalin penetration. Fixed specimens were transferred after at least 3 days from BSL3 facility to the Colorado State University Diagnostic Laboratories, BSL2 for trimming: Skulls were bisected (hemi skulls) and decalcified in semiconductor grade formic acid and EDTA (Calfor™, Cancer Diagnostics, USA) for 2-3 days. Oral cavity, salivary glands, olfactory bulb, cerebrum, cerebellum and brain stem were thoroughly inspected for gross lesions. Decalcified skulls and visceral organs were processed, embedded in paraffin wax and 4-5 µm sections were stained with hematoxylin and eosin for blinded evaluation by the pathologist using Nikon i80 microscope (Nikon Microscopy).

### Immunohistochemistry

Sections from hemi skulls and visceral organs were stained using ultraView universal alkaline phosphatase red detection kit. Heat-induced epitope retrieval was performed on a Leica Bond-III IHC automated stainer using Bond Epitiope Retrieval solution for 20 minutes. Viral nucleocapsid antigen was detected with a purified rabbit polyclonal antibody. Labeling was performed on an automated staining platform. Fast Red was used as chromogen and slides were counterstained with hematoxylin. Immunoreactions were visualized by a single pathologist in a blinded fashion. In all cases, normal and reactive mouse brain sections incubated with primary antibodies was used as a positive immunohistochemical control. Negative controls were incubated in diluent consisting of Tris-buffered saline with carrier protein and homologous nonimmune sera. All sequential steps of the immunostaining procedure were performed on negative controls following incubation.

### Immunofluorescence Staining and Tissue Imaging

Paraffin embedded tissue sections were stained for SARS-CoV-2 nucleocapsid protein (CSU; 1:500), microtubule-associated protein 2 (Abcam; ab32454; 1:500), ionized calcium binding adaptor molecule 1 (Abcam; ab5076; 1:50)/glial fibrillary acidic protein (Sigma; 3893;1:500) using a Leica Bond RXm automated staining instrument following permeabilization using 0.01% Triton X diluted in Tris-buffered saline (TBS). Blocking was performed with 1% donkey serum diluted in TBS. Sections were stained for DAPI (Sigma) and mounted on glass coverslips in ProLong Gold Antifade mounting medium and stored at ambient temperature until imaging. Images were captured using an Olympus BX63 fluorescence microscope equipped with a motorized stage and Hamamatsu ORCA-flash 4.0 LT CCD camera. Images were collected with Olympus Cellsens software (v 1.18) using an Olympus X-line apochromat 10X (0.40 N.A.), 20X (0.8 N.A.) or 40X (0.95 N.A.) air objectives, or Uplan Flour ×100 oil immersion (1.3 N.A.) objective.

### Next-generation sequencing library preparation for positive NP samples

Viral RNA from positive deer mouse samples was prepared for Next-generation sequencing. Briefly, cDNA was generated using SuperScript IV Reverse Transcriptase enzyme (Invitrogen) with random hexamers. PCR amplification was performed using ARTIC network V3 tiled amplicon primers in two separate reactions by Q5 High-fidelity polymerase (NEB). First-round PCR products were purified using Ampure XP beads (Beckman Coulter). Libraries were prepared using the Nextera XT Library Preparation Kit (Illumina) according to manufacturer protocol. Unique Nextera XT i7 and i5 indexes for each sample were incorporated for dual indexed libraries. Indexed libraries were again purified using Ampure XP bead (Beckman Coulter). Final libraries were pooled and analyzed for size distribution using the Agilent High Sensitivity D1000 Screen Tape on the Agilent Tapestation 2200, final quantification was performed using the NEBNext Library Quant Kit for Illumina (NEB) according to manufacture protocol. Libraries were then sequenced on the Illumina MiSeq V2 using 2 × 250 paired end reads.

### Deep sequencing analysis

Next-generation sequencing data were processed to generate consensus sequences for each viral sample. MiSeq reads were demultiplexed, quality checked by FASTQC, paired-end reads were processed to removed Illumina primers and quality trimmed with Cutadapt, duplicate reads were removed. Remaining reads were aligned to SARS-CoV-2 reference sequence by Bowtie2 (GenBank: MT020881.1). Alignments were further processed, and quality checked, using Geneious software, consensus sequences were determined and any gaps in sequences were filled in with the reference sequence. Consensus sequences were aligned in Geneious. Based upon the insertion that was identified in the passage 2 deer mice, a PCR assay was developed to detect the presence of the insert using a forward primer (5’-AGTGCGTGAT<u>AAGCTGAGAAGT</u>-3’; 12 nt insert sequence underlined) and reverse primer (5’-TAACCCACATAATAAGCTGCAGC) that generated a 183 bp product. A control forward primer (5’-GCACACGCCTATTAATTTAGTGC-3’) for detection of both wild-type (201 bp) or insert (213 bp) vRNA was designed 5’ to the insert primer.

### Model

The Phyre2 structure modeling engine (39) was used in the One-to-One Threading expert mode to generate a model of the deer mouse passage spike protein with the KLRS insert using the C chain of PDB entry 6VYB. The resulting model showed the insert is located at a surface loop on the N-terminal domain composed of residues 216-219 and predicts that this loop is enlarged to accommodate the 4-residue insertion. To illustrate the location of this insertion relative to the ACE2 receptor (Figure 4C), the structure of human ACE2 bound to the spike receptor binding domain (RBD), PDB entry 6VW1, was superposed on the up-conformation RBD from the B chain of the spike trimer.

## Acknowledgements

This work was supported by the Colorado State University Office of the Vice President for Research and by the National Institute of Allergy and Infectious Diseases (R01 AI140442). We thank Charles H. Calisher for constructive comments.

