## Supplemental Data for "SARS-CoV-2 infection, neuropathogenesis and transmission among deer mice: Implications for reverse zoonosis to New World rodents"

**Supplemental figure legends**

**Figure S1**. Serum response to SARS-CoV-2 structural proteins by infected deer mice. Protein from virus-enriched supernatants from SARS-CoV-2-infected Vero E6 cells, 8 µg protein per lane, were separated by SDS gel electrophoresis, and individual lanes from western blots were probed with 1:100 dilutions of sera from groups of three mock and infected deer mouse sera at times post infection as indicated and from contact controls at 28 dpi post exposure to infected animals. Prepared monoclonal antibody to the nucleocapsid protein served as control (N). Asterisks indicate predicted and known positions of structural proteins based on molecular weight: spike protein (S), cleaved spike 1 (S1), nucleocapsid (N), cleaved receptor binding domain (RBD), membrane (M), and envelope (E).

**Figure S2**. Schematic illustration depicting gustatory nerves carrying taste information from the soft palate and tongue to brain stem of the deer mouse. Chorda tympani nerve on anterior tongue (red) (1) and greater superficial petrosal nerve (green) (2) located on the soft palate along with conduct taste information from taste buds to cell bodies in the geniculate ganglion located within tympanic bulla. The glossopharyngeal nerve (CN IX, blue) conducts taste information from the circumvallate and posterior foliate papillae on posterior tongue to petrosal ganglion on the medial aspect of tympanic bulla. Information from taste ganglia is then transmitted into the nucleus tractus solitarius (NTS) (3) in medulla oblongata. In the medulla, taste responses are processed and then carried through parabrachial nucleus (PbN) and thalamic ventral posteromedial nucleus (VPM) to the primary gustatory nucleus in the insula. Perception of flavors is integrated with other avenues of sensation, particularly olfaction (1, 2). Temporary to persistent loss of taste is the one of the most consistent and common clinical symptoms reported in patients suffering from the mild form of COVID-19 (3). Pathologies of different components of the mouse gustatory system are following in Fig. S3A-I.

**Figure S3**. Necrosis of circumvallate and foliate papillae at posterior tongue, 3 dpi (A) with close-up view of neutrophils clustering around a taste bud (white arrow) (B). Dissecting edema dispersing lingual skeletal muscles apart, 3 dpi (C). Interstitial suppurative glossitis extending into branches of chorda tympani nerve (yellow arrow) at anterior tongue, 3 dpi (D). Multifocal viral immunostaining is showing lingual mucosa and submucosa with occasional immunoreactivity in a hypertrophied neuronal cell body in one of chorda tympani nerve branches, 6 dpi (E). Close-up view of the hypertrophied neuron in lingual parenchyma (F). Geniculate nucleus in tympanic bulla showing strong immunoreactivity in ganglionic neurons (cell bodies) of greater superficial petrosal and chorda tympani nerve fibers, 6 dpi (control negative –inset). Petrosal ganglion with attached CN IX shows strong immunoreactivity in corresponding ganglionic neurons 6 dpi. Strong immunoreactivity is seen in scattered neurons and glial cells mainly microglia in the area of the nucleus of the solitary tract (NST) (I). Bars = 100 µm.

**Figure S4**. Mouse olfactory (1, 4) and trigeminal pathways (2,3). The three major sensory branches of trigeminal nerve: ophthalmic (V1), maxillary, mainly formed by union between posterior nasal and rhinopalatine nerves (V2) and thickest branch, mandibular nerve (V3). Viral spread occurs via sympathetic and/or parasympathetic fibers into corresponding pterygopalatine and trigeminal ganglia (2 and 3, respectively). SARS-CoV-2 dissemination may be facilitated by transmission of disrupted olfactory epithelium and the olfactory neurons (1) to finally infect the main olfactory bulb (4) at 6 dpi as detailed below. Meningeal nerve was also inflamed at that time (4).

**Figure S5**. Massive necrosis and ulceration of MOE with expansion of submucosa by fibrinosuppurative exudate, 3 dpi (A). Close-up of embedded nerve branches, which appear rarefied by extensive status spongiosus and minimal infiltration of neutrophils, 3dpi (B). Maxillary sinuses from a control deer mouse show intact lining epithelium with no immunoreactivity (C), 3 dpi. Marked immunoreactivity is seen in lining and detached MOE, 3 dpi (D). Branches of facial nerve show moderate axonopathy (E). Pterygopalatine ganglion is multifocally cuffed by neutrophils with variable degeneration of the constituent neurons, 3 dpi (F). Ethmoidal nerves percolating cribriform plate showing multifocal suppurative neuritis and perineuritis (G). Severe congestion of meningeal vessels and rarefaction of meningeal nerve with histioneutrophilic perineuritis/meningitis , 3 dpi (H). Optic chiasm and hypothalamus show multifocal neuronal immunoreactivity, 6 dpi (I). Bars = 100 µm.

**Figure S6**. Retinal ganglionic cell bodies show multifocal immunoreactivity extending into inner plexiform layer and scattered bipolar cells in the inner nuclear layer, 3 dpi. Bar = 100 µm.

**Figure S7**. Calvarium bone marrow show strong cytoplasmic immunoreactivity in myeloid precursors, including monocytes and megakaryocytes, 6 dpi. Bar = 100 µm.

**Figure S8**. Grossly consolidated lung portions show massive infiltration of pulmonary parenchyma by numerous neutrophils and macrophages with peribronchiolar lymphoid hyperplasia (A). Main branches of pulmonary artery are infiltrated by neutrophils and cuffed by lymphoid follicles (B). Syncytial and histiocytic giant cells are dispersed among inflammatory cells expanding pulmonary interstitium and filling alveolar spaces (C). Lungs from a control mouse is within normal histologic limits with no immunoreactivity (D). Multifocal prominent bronchiolar and milder parenchymal immunostaining is seen in lungs 3 and 6 dpi (E). Mononuclear cells, macrophages, and antigen-presenting cells showing scattered cellular immunostaining in the paracortex (T-zone) area, 3 and 6 dpi (F). Bars = 100 µm.

**Figure S9**. Duodenum at 3 dpi shows mild focal histioneutrophilic duodenitis (A). Duodenum and ileum show moderate histioneutrophilic and lymphoplasmacytic enteritis (B and C). Small intestine from negative control deer mouse shows minimal nonspecific staining in goblet cells lining intestinal villi (D). Small intestine at 3 dpi shows marked immunoreactivity in the cytoplasm and apical border of mature enterocytes, crypt cells and scattered submucosal mononuclear cells consistent with macrophages (E). Bars = 100 µm.

**Figure S10**. Heart at 6 dpi, left subvalvular endocardium and myocardium shows minimal lymphocytic interstitial myocarditis (A) and more significant atrial suppurative perivascular myocarditis in right atrium (B). Kidney hilus shows mild perivascular suppurative inflammation that encircles small branches of renal nerves (C). Bars = 100 µm.

**Figure S11.** Detection of spike protein insert sequence by PCR. Forward primers were designed to amplify the wildtype or insert spike sequences. WT and insert were detected in both the deer mouse inoculum virus (Vero E6 passage 2) and original sourced BEI Resources virus. The BEI Resources virus had low amplification at 30 cycles that became greater at 40 cycles.

Table S1

Critical ACE2 residues of cricetid rodents involved in SARS-CoV-2 spike binding

| | | Susceptibility | ACE2 AA position | | | | | | | | | | | | | | | | | | | | ID | Accession |
|---|---|---|---|---|---|---|---|---|---|---|---|---|---|---|---|---|---|---|---|---|---|---|---|---|
| **Common name** | **Scientific name** | | 24 | 27 | 28 | 30 | 31 | 34 | 35 | 37 | 38 | 41 | 42 | 45 | 82 | 83 | 330 | 353 | 354 | 355 | 357 | 393 | | |
| **Human** | *Homo sapiens* | Yes | Q | T | F | D | K | H | E | E | D | Y | Q | L | M | Y | N | K | G | D | R | R | 20 | NP_001358344.1 |
| **Syrian hamster** | *Mesocricetus auratus* | Yes | Q | T | F | D | K | Q | E | E | D | Y | Q | L | N | Y | N | K | G | D | R | R | 18 | XP_005074266.1 |
| **Chinese hamster** | *Cricetulus griseus* | Unknown | Q | T | F | D | K | Q | E | E | D | Y | Q | L | N | Y | N | K | G | D | R | R | 18 | XP_003503283.1 |
| **Deer mouse** | *Peromyscus maniculatus* | Yes | Q | I | F | D | K | Q | E | E | D | Y | Q | L | N | Y | N | K | G | D | R | R | 17 | XP_006973269.1 |
| **White-footed mouse** | *Peromyscus leucopus* | Unknown | Q | I | F | D | K | Q | E | E | D | Y | Q | L | N | Y | N | K | G | D | R | R | 17 | XP_028743609.1 |
| **Dwarf hamster** | *Phodopus campbelli* | Unknown | Q | T | F | D | K | Q | E | E | D | Y | Q | L | N | Y | N | K | E | D | R | R | 17 | ACT66274.1 |
| **Prairie vole** | *Microtus ochrogaster* | Unknown | D | A | F | D | K | Q | E | E | D | Y | Q | L | S | Y | N | K | D | D | R | R | 15 | XP_005358818.1 |
| **Long-tailed pygmy rice rat** | *Oligoryzomys longicaudatus* | Unknown | V | T | F | D | N | Q | K | E | D | Y | Q | L | T | Y | N | K | G | D | R | R | 15 | PRJNA258076* |
| **Brown rat†** | *Rattus norvegicus* | No | K | S | F | N | K | Q | E | E | D | Y | Q | L | N | F | N | H | G | D | R | R | 13 | NP_001012006.1 |
| **House mouse†** | *Mus musculus* | No | N | T | F | N | N | Q | E | E | D | Y | Q | L | S | F | N | H | G | D | R | R | 13 | NP_001123985.1 |

* See Campbell et al., 2015 (PMID 25856432)
† Muridae rodents

Table S2. Primers used for SYBR Green qPCR gene expression profiling (5'→3')

| Gene | Forward | Reverse |
|---|---|---|
| Gapdh | GGTGCCAAAAGGGTCATCATCTC | GCAGGAAGCGTTGCTGATAATCTTG |
| Foxp3 | AAGCAGATCACCTCCTGGATGAG | TAGCACCCAGCTTCTCCTTTTCC |
| Gata3 | AGTCCGCATCTCTTCACCTTCC | GGCACTCTTTCTCATCTTGCCG |
| Tbx21 | GCCAACCCCAAGGATATGGT | GAATGTGGGCTTCATGCTC |
| CD4 | GGTTGAAATGAAGGACCCTGAGG | CCTCTGGATGAAACCTGGATTTTGG |
| CD8a | AAGAAGAGCGGATTGGACTTCG | AGATGAGAGTGATGACCAGGGACAG |
| TCRb | CAATAACATCACCTACTGCCTGAGC | GTAAGCCCATAGAACTGGACTTGG |
| TGFb | CGTGGAACTCTACCAGAAATACAGC | TCAAAAGACAACCACTCAGGCG |
| Ifng | GGCTATTTCTGGCTGTTACTGCC | ATCCCCGACATCTGAGCTACTTG |
| Il4 | CCCCGTGCTTGAAGAACAATTC | GGACTCATTCCCAGTACAGCTTTTC |
| Il4ra | AGAACCCTGTTCCCAACCA | TTGGATGGCAACTCCATGT |
| Il13 | TGCAAACCCATCTACAAGACCC | GCCACTTCGATTTTGGTATCCG |
| Il13ra1 | TGGTGTTCTTCCTGATGCTG | AGCGCTTCGCTCCAATTA |
| IL21 | AACTCAAGCCAGCAAACACAGG | GCTGCTTTTTCTCAGCCTTGG |
| IL23 | AGAAATGATGTCCCCCGTATCC | CAGACCTTGGTGGATTCTTTGC |
| Il2ra | TGCCACATTCAAAGCCCT | CCAACTCCTTTGTTCTTCGG |
| Socs1 | TGAGATCGCGAAGAACCTG | GGAAGGGGAAGGAACTTCA |
| Socs5 | CGGTTTGGGGACCATTTTA | TCAATCCCCGTTCTGCAT |
| Tnf | TGTAGCCCACGTTGTAGCAAACC | CTGGTTGTCTTTGAGATCCATGC |
| Zeb1 | CCCAACAAACAGCTGCAA | ATCCACAGCCCCATCAAA |
| CD80 | CATTGTGTGTCGCGTTGTGT | TGTCGGGGTCACCTCAGTTA |
| CD86 | CTTACCTTGCCAGCTCTGCT | TGCCCTAGACCTGAGTGTGA |
| Ctla4 | CTTGTCTTGGAGTCCCGAGG | ATGGCTTTGGAGAAGGTCGG |
| Ccl2 | CAGACGTACACAAGAAAACTGGACC | GTCAAGTTCGCATTCAAAGGTGC |
| Ccl3 | AGCCAGGTGTCATTTTCCTAACC | CAGCTCAGTGATGTATTCTTGGACC |
| Ccl5 | CCACGTCAAGGAGTATTTCTACACC | TCCTGAACCCACTTCTTCTTTGG |
| Cxcl10 | CACTGTCAGCACCATGAACC | GGGATTCCTTGAGTCCCACT |
| IFNg | GGCTATTTCTGGCTGTTACTGCC | ATCCCCGACATCTGAGCTACTTG |
| TGFb | CGTGGAACTCTACCAGAAATACAGC | TCAAAAGACAACCACTCAGGCG |
| Il10 | TAAGGGTTACCTGGGTTGCCAAG | CAAATGCTCCTTGATTTCTGGGC |
| Il12a | TCCAAAAACCTGCTGAGGACCAC | AATCAACGTCTTCAGGAGTGCAG |
| Il12b | TGTTCTCATGGGCTGATCC | GGCAGCCTTGGTTGAAAA |
| Il6 | CCATCCAACTCATCCTGAAAGC | CCACAGATTGGTACACATAGGCAC |
| Ifna2 | ACTCATCTGCTGCTTGGGA | TTGCAGGTCTTTGAGCTGG |
| Ifnb1 | TGCCTTCGTCATCCAAGAG | TCTCATTCCACCCAGTGCT |
| Irf3 | TAGAAAAGGAAGCCCCAGC | TGGTGCCAACACTGGTTTC |
| Isg15 | AGAGTTCCTGGTGTCCCTGA | ACACCAGTCTTCTGGGCAAT |
| Mavs | GCAACTAGGGAACCAGGACA | AAGCACTGTACCCAGCCAAC |
| Tbk1 | TCTCGAATAGCCAGCACCTT | CTCACCAGCTCAATCAACCA |
| TLR7 | CCTGGTTGCCTGAATCTTTC | TGAGCAATGGCAGCTGTTAG |
| Oas2 | TGGACGAGTTCGACATTGC | AAGATGCAGAGCTGCTGGT |
| Pycard | ACCAGCACAGACAAGCACTG | GTCAGCACACTGCCAAACAG |

Figure S1.

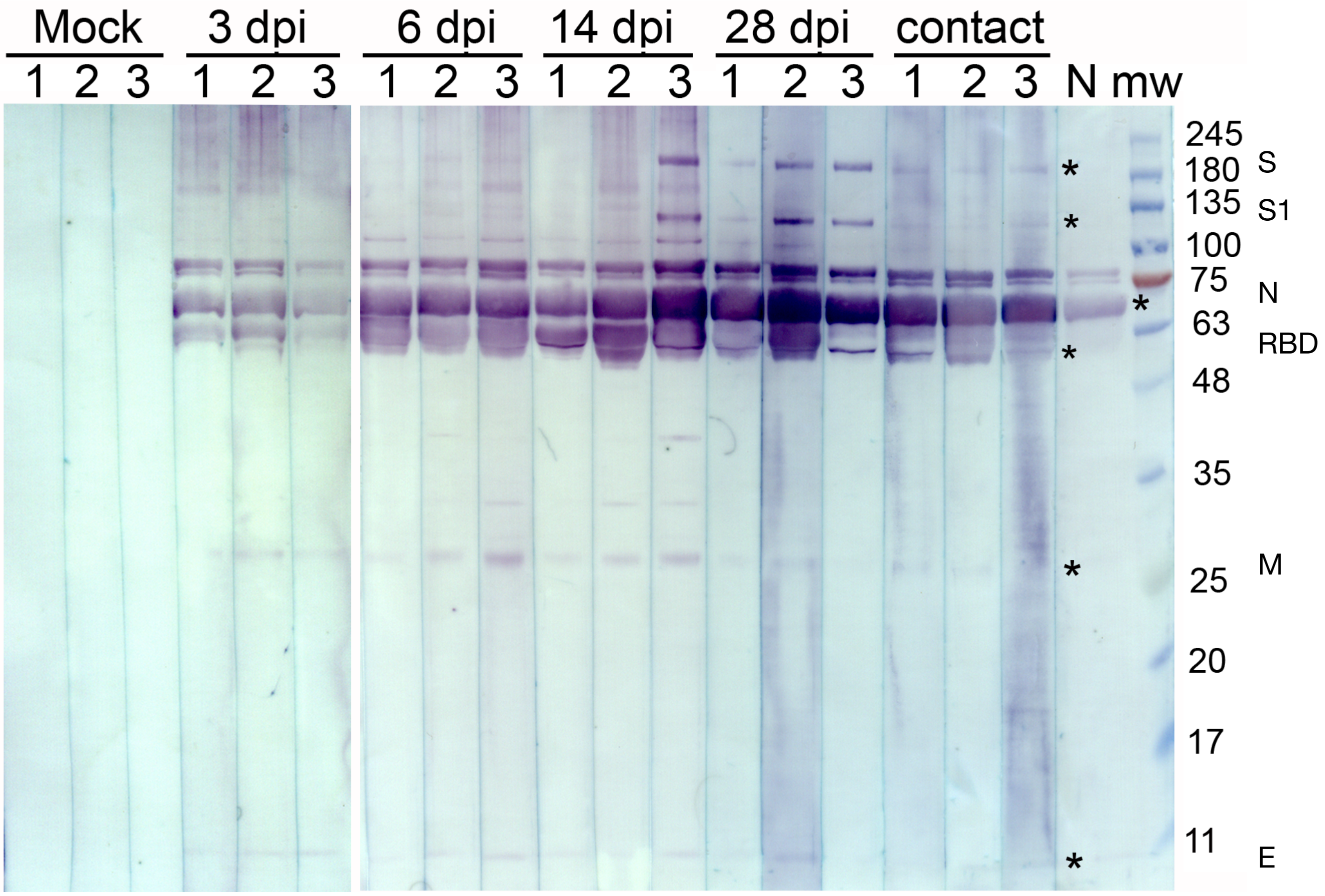
Mock
3 dpi
6 dpi
14 dpi
28 dpi
contact
1 2 3 1 2 3 1 2 3 1 2 3 1 2 3 1 2 3 N mw
245
180
135
100
75
63
48
35
25
20
17
11
S
S1
N
RBD
M
E

Figure S2

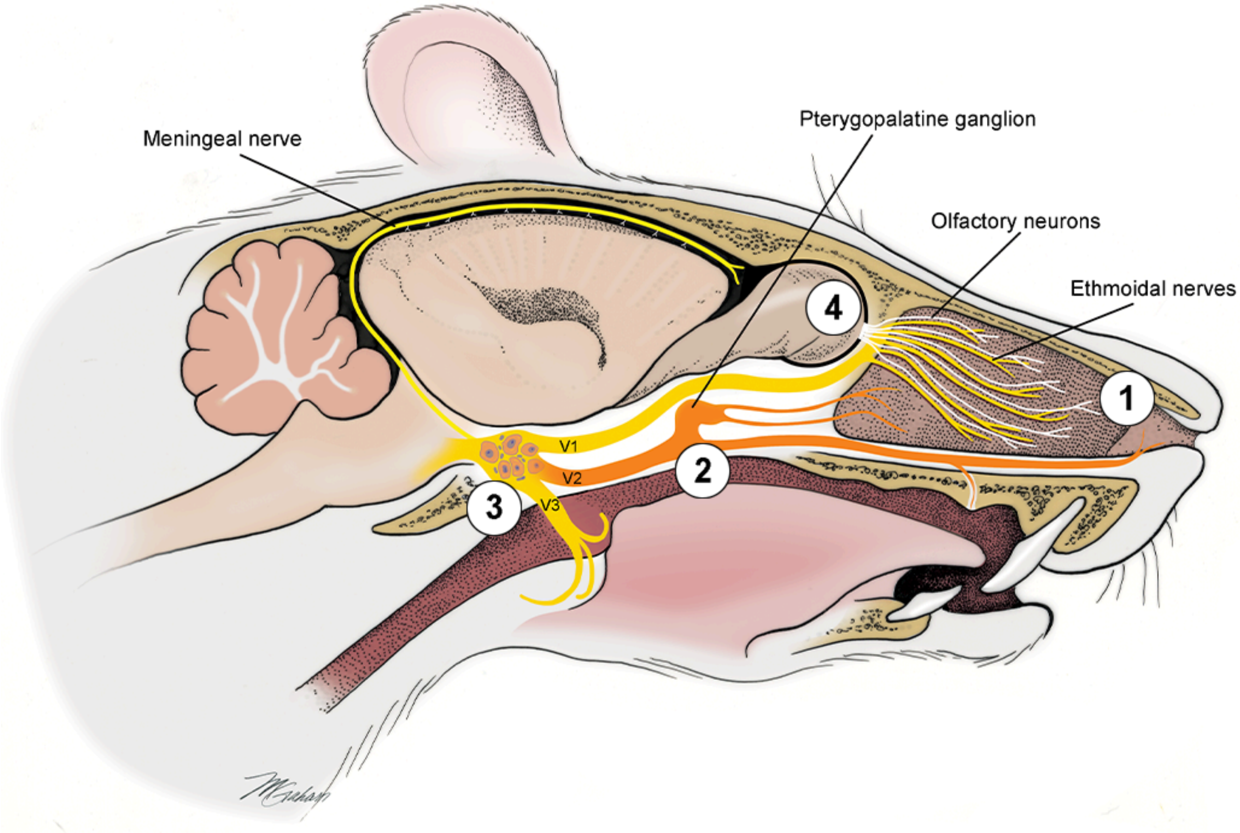
Meningeal nerve
Pterygopalatine ganglion
Olfactory neurons
Ethmoidal nerves
4
1
V1
V2
V3
2
3

Figure S3

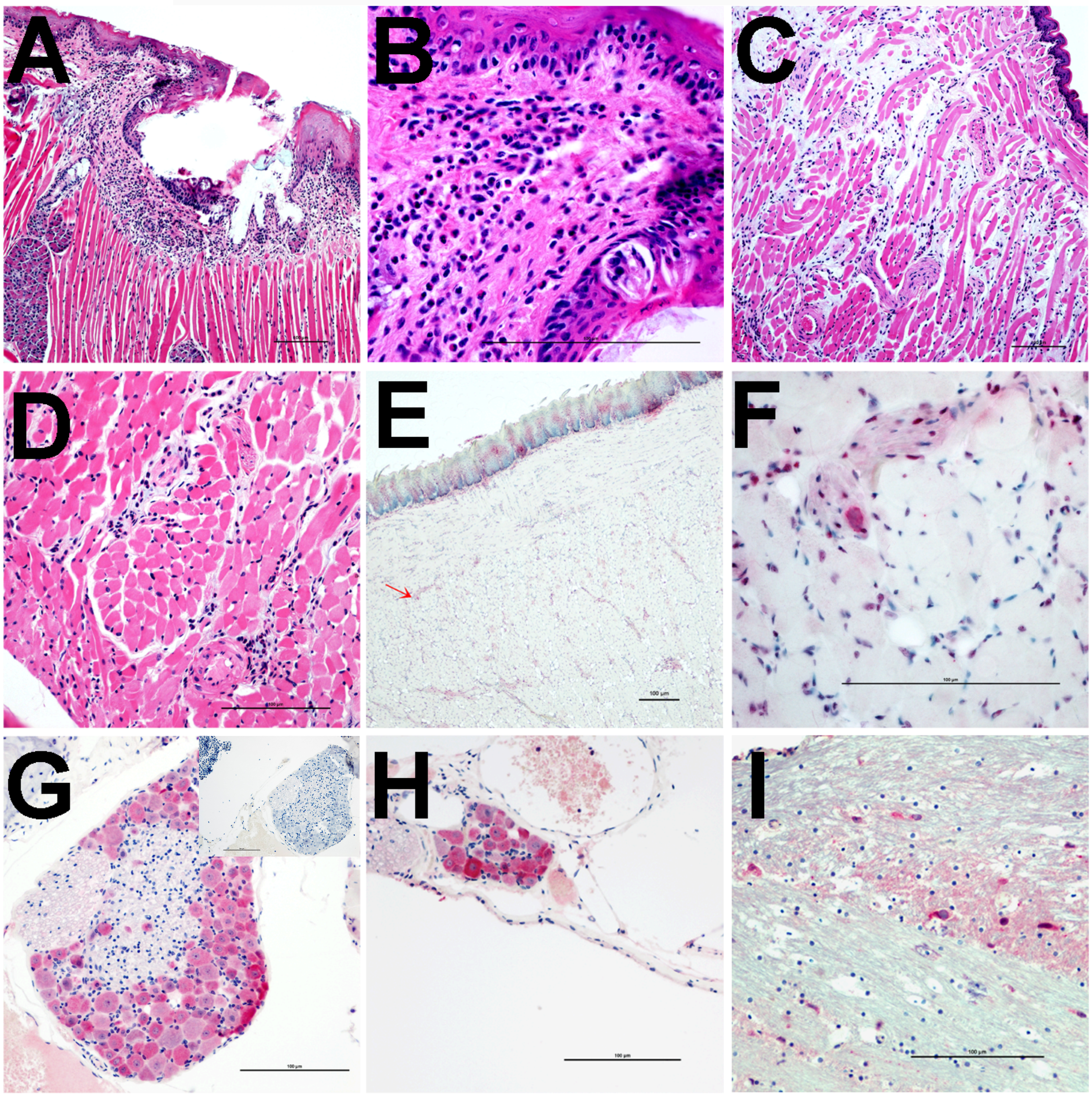

Figure S4

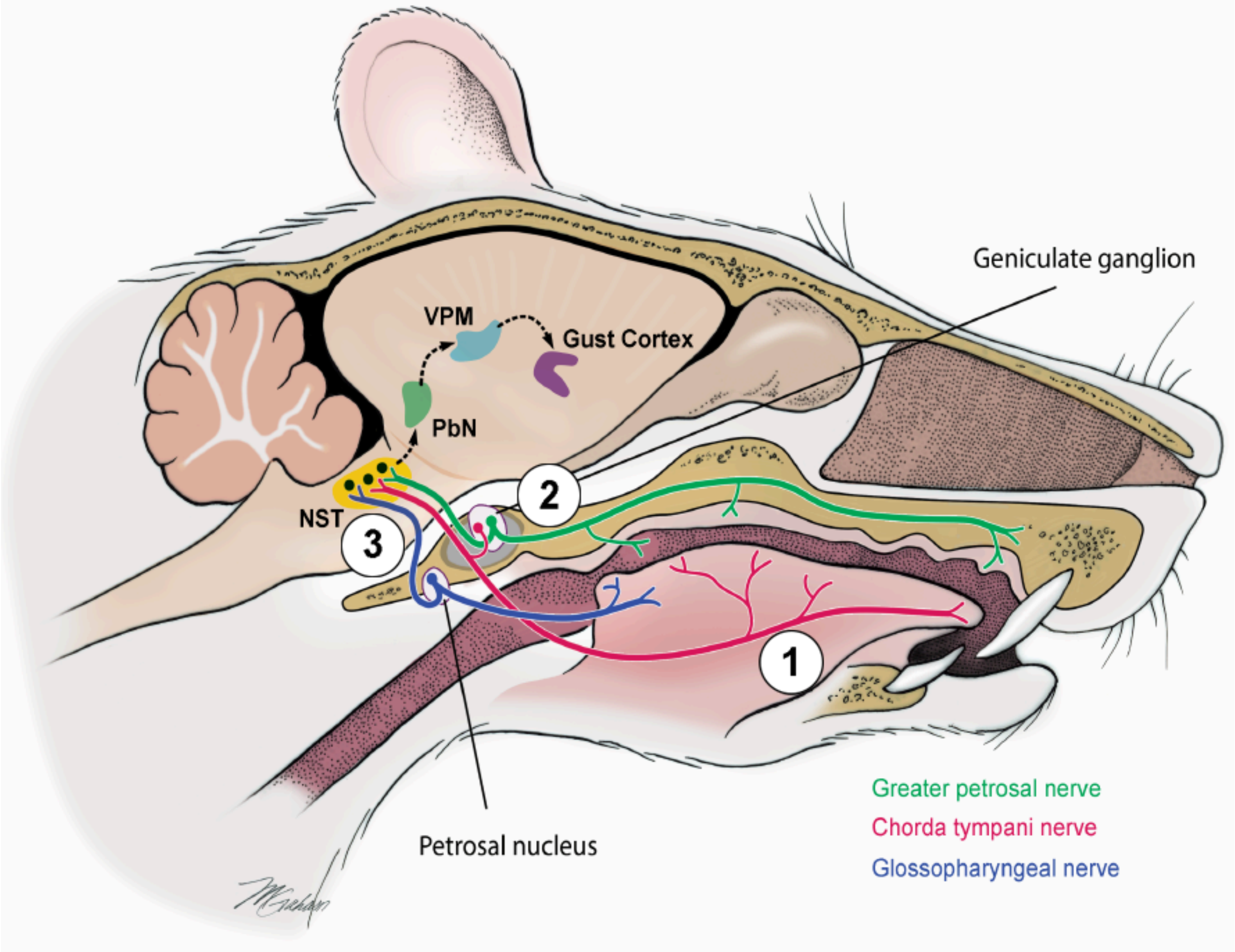
Geniculate ganglion
VPM
Gust Cortex
PbN
NST
2
3
1
Petrosal nucleus
Greater petrosal nerve
Chorda tympani nerve
Glossopharyngeal nerve

Figure S5

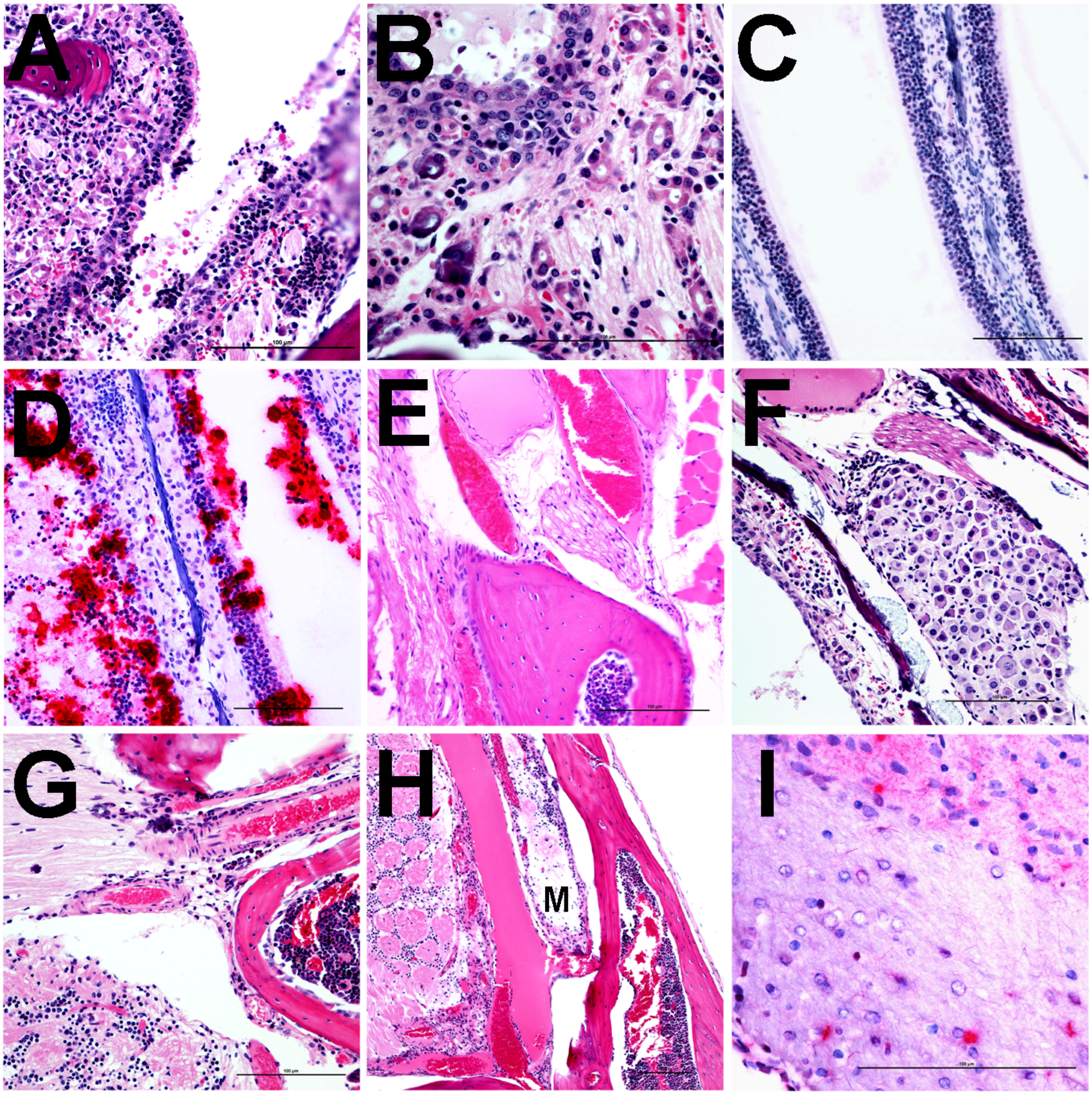

Figure S6

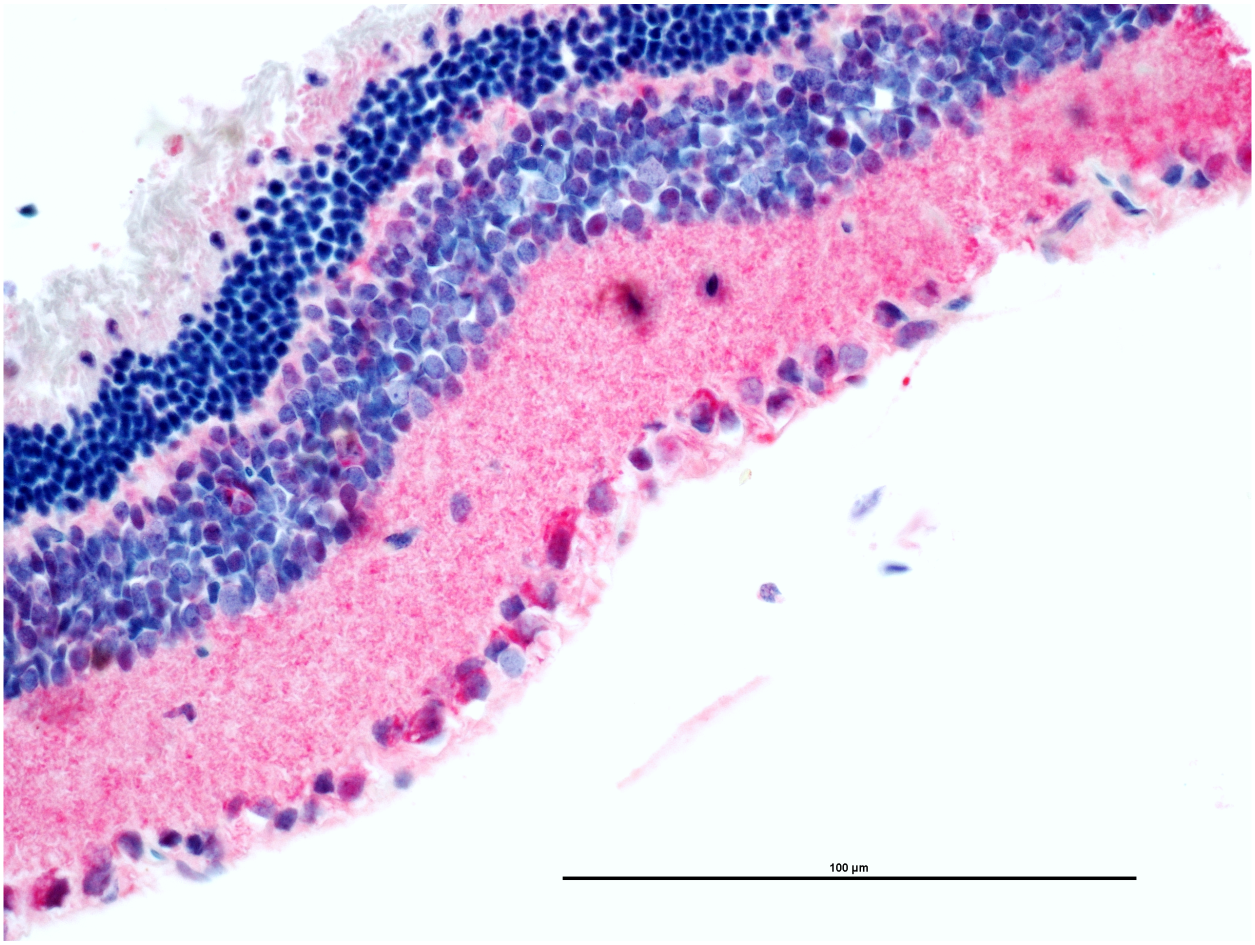

Figure S7

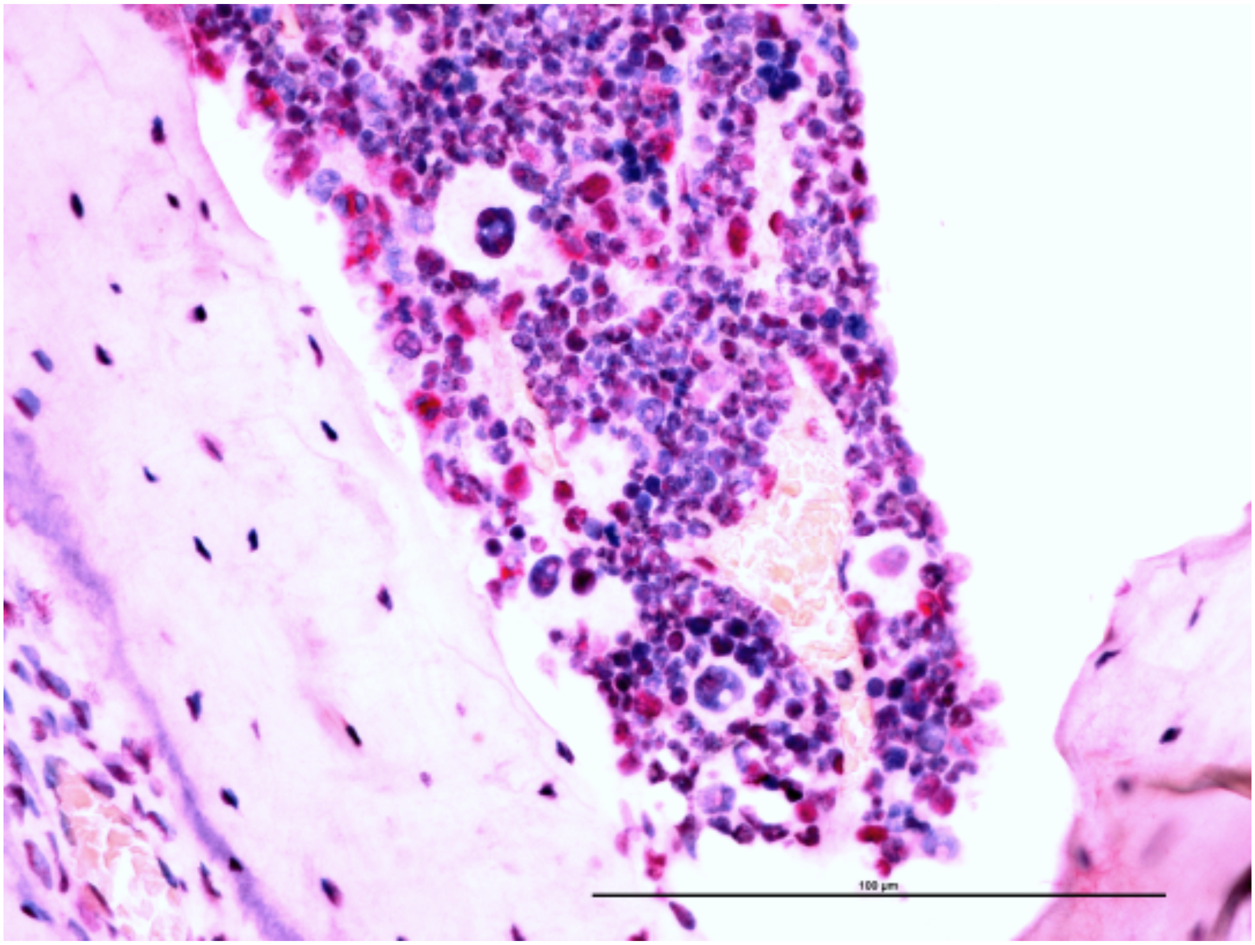

Figure S8

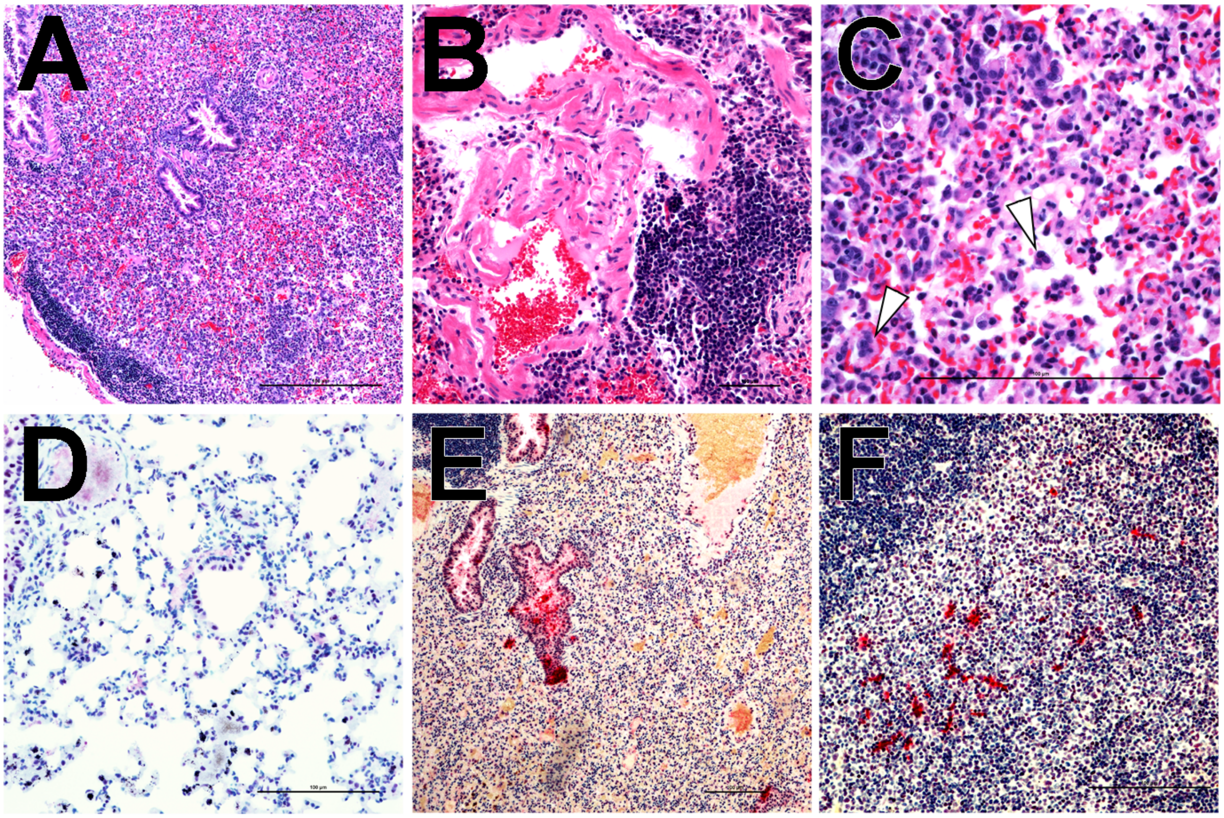

Figure S9

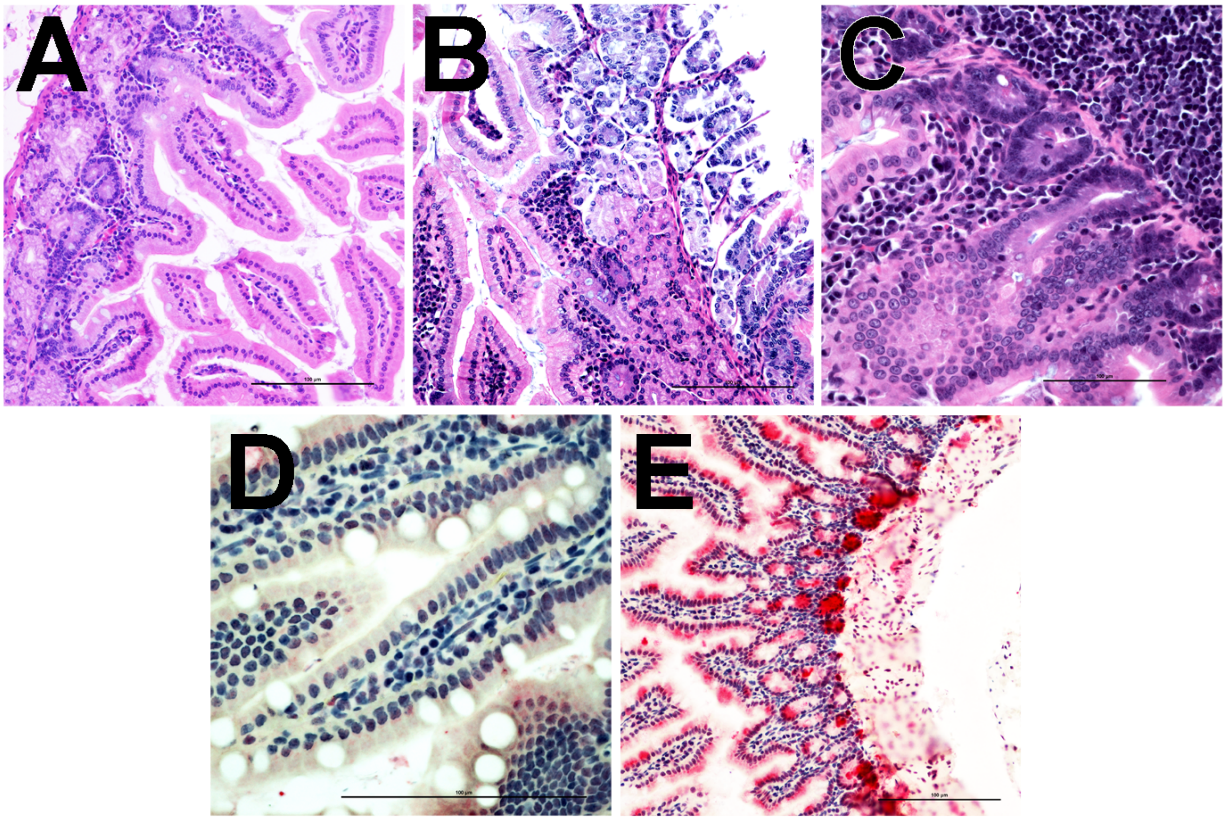

Figure S10

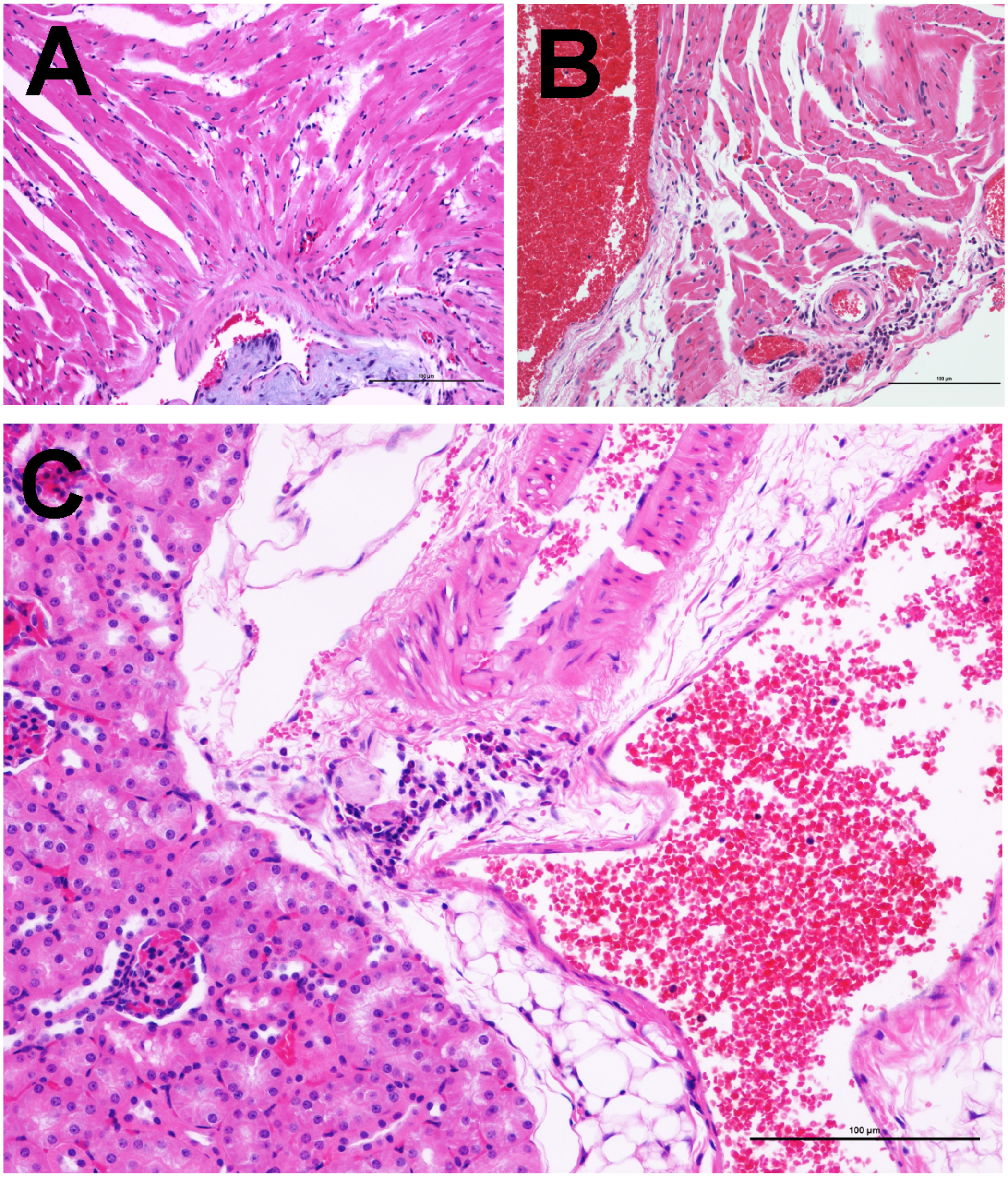

Figure S11

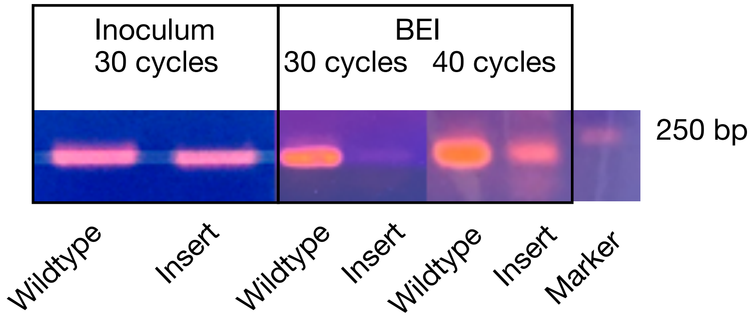
Inoculum
30 cycles
BEI
30 cycles
40 cycles
250 bp
Wildtype
Insert
Wildtype
Insert
Wildtype
Insert
Marker
